# Enhancement of larval immune system traits as a correlated response to selection for rapid development in *Drosophila melanogaster*

**DOI:** 10.1101/029439

**Authors:** Punyatirtha Dey, Kanika Mendiratta, Joy Bose, Amitabh Joshi

**Affiliations:** Evolutionary Biology Laboratory, Evolutionary & Organismal Biology Unit, Jawaharlal Nehru Centre for Advanced Scientific Research, Jakkur P.O., Bengaluru 560 064, India.

**Keywords:** developmental rate, hemocytes, phenol oxidase, life-history tradeoff

## Abstract

We have shown earlier that the evolution of rapid development is accompanied by a correlated decrease in larval feeding rate and competitive ability in laboratory populations of *Drosophila melanogaster* (Prasad et al. 2001; Shakarad et al. 2005). Here, we show that our faster developing populations have evolved higher hemocyte density and phenol oxidase activity in the larval hemolymph. The increased hemocyte density could be responsible for the evolution of decreased feeding rate as hemocytes and the cephalopharyngeal musculature share common embryonic precursor cells (Kraajiveld et al. 2001). We also show that the bacterial load in larval food vials of the faster developing populations is substantially higher than in controls. Our results suggest that the evolution of reduced competitive ability in the faster developing populations is probably due to larval feeding rate trading off with enhanced larval immune system function. Enhanced larval immune function, in turn, is most likely selected for due to the role of hemocytes (Lanot et al. 2001, Wood and Jacinto 2007) and phenol oxidase (Pentz et al. 1986) in development, and perhaps also due to inadvertent selection on immune performance resulting from the higher bacterial load faced by larvae in the faster developing populations.

## Introduction

In nature, *Drosophila* larvae typically inhabit ephemeral habitats like rotten fruits, often at high density (Atkinson and Shorrocks 1997), conditions that are believed to favour the evolution of rapid pre-adult development (Clarke et al. 1961; Partridge and Fowler 1992). However, rapid development is also correlated with reduced adult body size (Chippindale et al. 1997; Nunney 1996), which is itself a strong correlate of reproductive fitness. Development time, thus, is a key life-history trait linking fitness components in the pre-adult and adult life-stages, and the various tradeoffs surrounding development time are believed to mediate the evolution of developmental and growth rates and body size in *Drosophila*. Consequently, direct and correlated responses to selection for rapid pre-adult development have been extensively studied in *D. melanogaster* over the last two decades or so, yielding important insights into the fitness consequences of genetic constraints that shape the evolution of development time (reviewed by Prasad and Joshi 2003).

For example, two sets of populations of *D. melanogaster* selected for rapid development in our laboratory independently evolved reduced larval feeding (cephalopharyngeal sclerite retraction) rate (Prasad et al. 2001; Rajamani et al. 2006). Most likely due in part to the reduced feeding rate, larvae from the faster developing populations were poorer competitors at high larval rearing density than their ancestral control counterparts (Joshi et al. 2001; Shakarad et al. 2005). This tradeoff between developmental rate and feeding rate indicates possible competition between these two traits for some common underlying resources. However, nothing is presently known about the nature of the limiting resources that affect the expression of both feeding rate and development rate, although studies on parasitoid-resistant populations of *D. melanogaster* suggest a possible explanation.

The evolution of increased parasitoid resistance in *D. melanogaster* under laboratory selection is accompanied by the correlated evolution of lower competitive ability due to a reduced larval feeding rate (Fellows et al. 1999; Kraajiveld et al. 1997), and an increased hemocyte density in the larval hemolymph (Kraajiveld et al. 2001). Upon parasitoid attack, larvae from these parasitoid resistant populations mount a more vigorous hemocyte-mediated encapsulation response compared to relatively susceptible controls (Fellows et al. 1999; Kraajiveld et al. 1997, 2001). Since both the hemocytes and the cephalopharyngeal musculature of larvae originate in the head mesoderm of the embryo, it appears that the parasitoid resistant larvae invest limiting cellular resources into hemocytes rather than cephalopharyngeal musculature during development and, consequently, have a reduced larval feeding rate that ultimately translates into reduced competitive ability (Kraajiveld et al. 2001).

While hemocytes are known to be important for defense against pathogens/parasitoid eggs in *Drosophila* (Lemaitre 2007), they are also essential for proper development (Lanot et al. 2001, Wood and Jacinto 2007). During metamorphosis, the hemocytes engulf the cells marked for destruction, thus playing a major role in tissue remodeling (Lanot et al. 2001). Hemocyte-deficient mutants of *Drosophila* do not complete metamorphosis and die as prepupae (Braun *et al*. 1998). Given the link between higher hemocyte density and decreased larval feeding rate (Kraajiveld et al. 2001) on the one hand, and hemocytes and development on the other, we hypothesized that our faster developing populations may have evolved a higher hemocyte density to be able to support metamorphosis in a substantially reduced pupal duration, about three-fourths that of controls (Modak 2009). The necessity of rapid development in these populations could therefore be responsible for the correlated evolution of reduced larval feeding rate.

The hemolymph of *Drosophila* larvae contains three types of hemocytes, namely plasmatocytes, crystal cells and lamellocytes (Braun et al. 1999). Plasmatocytes act as phagocytes, while crystal cells store phenol oxidase and other enzymes required for the melanization response (Braun et al. 1999, Lemaitre and Hoffman 2007). Lamellocytes are normally absent from hemolymph, and differentiate only on infection by parasitoids (Lemaitre and Hoffman 2007). In this study, we estimated the hemocyte density and phenol oxidase activity in larvae of our fast developing populations of *D. melanogaster*. We also investigated whether there was any difference between faster developing populations and controls in the bacterial load in the larval food medium of their respective culture vials.

## Materials and methods

### Experimental populations

We studied eight populations of *D. melanogaster*, of which four were ancestral controls (JB_1-4_) and four had been subjected to selection for rapid development and early reproduction (FEJ_1-4_) for about 350 generations at the time of this study. These populations and their maintenance regimes have been described in detail earlier (Prasad et al. 2000, 2001). Briefly, each FEJ population was derived from a single JB population: FEJ*_i_* derived from JB*_i_*, *i* = 1-4. All populations were maintained on a discrete generation cycle, at 25 ± 1°C and constant light and high humidity, on banana-jaggery food medium. The number of breeding adults was about 1800 in the JBs and about 1200 in the FEJs. The adult flies were maintained in Plexiglas cages (25 × 20 × 15 cm^3^), and the larvae in 8-dram glass vials (9 cm height × 2.4 cm diameter). The JB controls were maintained on a 21 day discrete cycle, with the eclosed adults being transferred to fresh vials on the 12^th^, 14^th^ and 16^th^ day after egg collection. Twelve days are sufficient for all viable individuals to eclose at the rearing larval density of 60-80 eggs per 6 mL of food per vial used. On the 18^th^ day after egg collection, the flies were collected into a Plexiglas cage containing a Petri dish of food covered with yeast-acetic acid paste. The eggs laid by these flies on the 21^st^ day were collected off a fresh Petri dish with food and transferred to vials. The maintenance regime of the FEJs differed from the JBs in two important aspects, together imposing selection for rapid development and early reproduction relative to the controls. First, only the earliest 25 percent or so of the flies that eclosed in each food vial were collected into cages to form the breeding adult population. Second, the eggs from these adult flies were collected 3 days post-eclosion to initiate the next generation. At the time this study, the mean egg-to-adult development time for the FEJs was 7.1 days, compared to 9.3 days for the JBs (Modak 2009).

To eliminate any non-genetic parental effects of the different maintenance regimes, all eight JB and FEJ populations were maintained under identical control conditions for one generation prior to assays. Eggs collected from these “commongarden” flies were used to generate larvae, which were then used for the assays. Late 3^rd^ instar wandering stage larvae were used for all the assays.

### Measurement of hemocyte density in larval hemolymph

Thin glass capillaries were pulled in flame to generate ultra-thin glass needles, which were then attached to 10 *µ*L pipette tips. Next, five larvae from a single population were placed on a slide, and their posterior ends were cut with scissors. The hemolymph from the five larvae was pooled on the slide. Looking through a stereomicroscope, 0.5 *µ*L of hemolymph was pipetted out, diluted with 0.05 M phosphate buffer saline, pH 7.5, and added to the counting chamber of a hemocytometer (Rohem Instruments, Nasik, India). The number of hemocytes was manually counted at 400X using an Eclipse E 200 microscope (Nikon, Guangdong, China). This procedure was carried out on larvae from all eight FEJ and JB populations, with five independent replicates of five pooled larvae each per population.

### Measurement of phenol oxidase activity per µL of larval hemolymph

As described above, 5-10 larvae were bled on a glass slide and 1 *µ*L of hemolymph was pipetted into 7 *µ*L of 10 mM DOPA in 10 mM PBS, pH 6.5, and activated for 10 minutes at room temperature. After every 5 minutes, 1 *µ*L of reaction mixture was taken and the optical density at 470 nm was measured in a Nanodrop 1000 spectrophotometer (Thermo Scientific, Wilmington, USA). Change in optical density at 470 nm per hour was calculated from the linear region of the reaction, corrected for the dilution factor, and designated as phenol oxidase (PO) activity. For each population, five independent replicates were run and PO activity per *µ*L of hemolymph was calculated for each replicate.

### Measurement of bacterial density in the larval food medium

For each population, 70 eggs were placed on 6 mL of banana food in each of 10 replicate vials, and the larvae were allowed to develop for 3 days. After 3 days, 20 mL Luria broth (LB) was added to each vial and the food was lightly homogenized with a glass rod and suspended in LB. The suspension was centrifuged at 100 g for 5 minutes and the supernatant was diluted and plated on LB agar plates. The plates were incubated at 37ºC for 18 hours, and colonies were counted manually to determine the bacterial density per mL of larval food for each vial.

### Statistical analyses

The independent measurements of total hemocyte count, PO activity and bacterial density were taken as replicates within each population, and the data subjected to separate mixed model analyses of variance (ANOVA), with block, comprising of one FEJ population and the corresponding ancestral JB population, treated as a random factor crossed with selection regime. All analyses were implemented on Statistica^TM^ for Windows Release 5B (Statsoft Inc. 1995).

## Results

### Hemocyte density in larval hemolymph

The mean hemocyte density in the hemolymph of FEJ larvae was found to be 3.9 fold more than that in the ancestral control JB populations (Table 1), and the difference was significant (*F*_1,3_ = 78.028, *p* = 0.0030).

**Table 1.** Mean (± s.e.) immune system related traits and bacterial loads in control (JB) and rapidly developing (FEJ) populations. Standard errors reflect the variation among means of the four replicate populations within each selection regime.

| Selection Regime | Hemocyte density | PO activity | Density of non pathogenic bacteria in food |
| --- | --- | --- | --- |
| | (Number of hemocytes per mm<sup>3</sup> larval hemolymph) | (ΔOD<sub>470</sub> per μL of hemolymph per hour) | (Colony forming unit $\times 10^{-3}$/mL) |
| JB | 5794 ± 511 | 3.92 ± 0.24 | 6.0 ± 1.9 |
| FEJ | 22685 ± 2052 | 15.04 ± 0.81 | 35.6 ± 4.2 |

### Phenol oxidase activity per unit volume of larval hemolymph

The mean PO activity per *µ*L hemolymph in FEJ larvae was found to be a 3.8 fold more than the ancestral control JB populations (Table 1), and the difference was significant (*F*_1,3_ = 282.492, *p* = 0.0005).

### Bacterial load in larval food medium

The amount of bacteria present in the larval food vials of the FEJ populations was found to be 5.8 fold more than that in the ancestral control JB populations (Table 1), and this difference was significant (*F*_1,3_ = 47.49, *p* = 0.0063)

## Discussion

Our observation that the faster developing FEJ populations, earlier seen to have evolved decreased larval feeding rate (Prasad et al. 2001) and competitive ability (Shakarad et al. 2005), have also evolved a higher hemocyte density than ancestral controls (Table 1) is consistent with the tradeoff between hemocyte density and feeding rate of larvae from parasitoid resistant populations of *D. melanogaster* observed by Kraajiveld et al. (2001). Our finding thus supports the notion of competition for progenitor cells between investment in cephalopharyngeal musculature and hemocyte levels during early development (Kraajiveld et al. 2001). Accordingly, the correlated evolution of reduced larval feeding rate in the FEJ populations is likely to be driven at least in part by the necessity of increased hemocyte density for rapid development.

We also found that the FEJ larvae had increased PO activity per unit volume of hemolymph, compared to the ancestral JB controls (Table 1). PO activity in hemolymph is considered an immune parameter in insects (Lemaitre 2007) and is also related to the darkening of pupal and adult cuticle during development (Pentz et al. 1986). Overall, our finding that both hemocyte density and PO activity in the larval hemolymph increased in the FEJs also provides strong support for the notion of pleiotropy between developmental and immune functions in *Drosophila* larvae (Yixin et al. 2009). Yixin et al. (2009) selected populations of *D. melanogaster* for larval resistance to the pathogen *Pseudomonas aeruginosa* and observed the correlated evolution of reduced development time in the resistant populations. Our results suggest that the response of development time and larval immune system function to selection on one or the other trait is symmetric, a finding that strengthens the inference of pleiotropy between these traits.

Due to the dual role of hemocytes and PO in development and immunity, it is however difficult to infer unequivocally whether these traits in the FEJ populations evolved to facilitate faster development or in response to increased load of possibly nonpathogenic bacteria in the food medium, or both. A recent study on the insect *Trichoplusia* showed that even non-pathogenic, non-infectious bacteria present in the food of larvae can elicit an immune response and affect fitness components like pupation time (Freitak et al. 2007). In *Drosophila* cultures, microbial growth is suppressed by the feeding activity of larvae (Sang et al. 1949) and cultures with very few larvae per vial often get overgrown with bacteria and yeast compared to cultures with several tens of larvae per vial (A. Joshi, P. Dey, *pers. obs.*). Given that third instar FEJ larvae are smaller and slower feeders than their control counterparts but are maintained at the same density per vial (Prasad et al. 2001), it is possible that their inhibitory effect on bacterial growth in the larval food vials is less than that of the JB controls. The nature of this inhibitory effect of *Drosophila* larvae on microbial growth in the food vials is not clear. It could be that larvae check microbial population growth merely by feeding, or that larvae inhibit microbial growth through some hitherto unknown mechanism. Whatever the mechanism, however, it is clear that the FEJ culture vials harbor substantially larger bacterial populations at the third instar stage than control vials. In principle, this increased bacterial load could also directly select for enhanced immune system performance in the FEJ populations, independent of the effect of selection for rapid development, mediated through the multi-tasking role of hemocytes and melanization. Such inadvertent selection pressures are not unusual in experimental evolution studies (Prasad and Joshi 2003).

We have earlier found that our FEJ populations suffer greater adult mortality than controls when grown in the presence of *Escherichia coli* (Modak et al. 2009). Unlike larval immune function, adult immune function is seen to trade off with development time in insects: populations of the yellow fever mosquito, *Aedes aegypti*, selected for rapid development showed reduced adult immunity (Koella and Boete 2002) whereas selection for higher adult immunity in the Indian meal moth, *Plodia interpunctella*, resulted in correlated evolution of longer development time (Boots and Begon 1993). In general, it appears that larval and adult immunity in *Drosophila* have the potential to be correlated or uncorrelated, depending on which measures of immune function are assayed (Petersen et al. 1999; Fellous and Lazzaro 2011). At this time, it is difficult to say whether our present results together with those of Modek et al. (2009) indicate a specific tradeoff between larval and adult immune function in our populations. As noted by Modak et al. (2009), it is possible that the increased mortality of the FEJ adults in the presence of *E. coli* is due to their mounting an equal or greater immune response than JB adults and then succumbing faster to the starving conditions of the assay vials as they have less lipid reserves to start with. What does seem clear, however, is that the FEJ larvae exhibit increased immune function, at least for some measures of larval immunity.

## Acknowledgments

We thank N. Rajanna and M. Manjesh for assistance in the laboratory. PD was supported by a Research Associateship from the Jawaharlal Nehru Centre for Advanced Scientific Research (JNCASR). KM was supported by a fellowship through the Project Oriented Biology Education program of the JNCASR. The work was supported in part by funds from the Department of Science and Technology, Government of India. Preparation of the manuscript was supported in part by a J. C. Bose National fellowship from the Department of Science and Technology, Government of India, to AJ.

